## Supplemental Information for "CNAViz: An interactive webtool for user-guided segmentation of tumor DNA sequencing data"

### A CNAViz Features

#### A.1 Data Characteristics

CNAViz input data adhere to seven characteristics.

*(D1) One or more samples, quantified by* $m > 0$*, are sequenced from a tumor.* Samples may correspond to bulk DNA sequencing samples and/or single-cell DNA sequencing samples.

*(D2) The genome is partitioned in* $n$ *bins that may vary in size.* We indicate the chromosome in which bin $i$ occurs by $\mathrm{chr}(i)$, its start position on that chromosome by $\mathrm{start}(i)$ and end position by $\mathrm{end}(i)$.

*(D3) The read depth ratio* $\mathrm{RDR}(p, i)$ *is provided for each bin* $i$ *in each sample* $p$*.*

*(D4) The B-allele frequency* $\mathrm{BAF}(p, i)$ *is provided for each bin* $i$ *in each sample* $p$*.*

*(D5) Optionally, each bin may be assigned to a segment/cluster* $\mathrm{cluster}(i)$. These values represent the local or global segmentation performed by existing algorithms and is used for further refinement.

*(D6) Optionally, a set* $D$ *of driver genes may be provided with genomic coordinates* $(\mathrm{chr}(d), \mathrm{start}(d), \mathrm{end}(d))$ *for each driver gene* $d$*.*

*(D7) Export new clustering.* The new clustering created is exportable for future usage. Bins that have been erased (C5) are excluded from export.

#### A.2 Usability

*(U1) Color dropper*. Allows the user to choose the color of the selected bins. The default color is black.

*(U2) Point size slider*. Allows the user to change the size of the point representing each bin in all the plots.

*(U3) Tooltips*. Hovering over each button will provide the user with a description of that button.

*(U4) Toolbar*. The toolbar floats at the top of the screen, and can be used to switch between the four different modes (Zoom, Pan, Select, Erase).

*(U5) Help*. A help button is located in the top right corner.

#### A.3 Analysis Tasks

**Plotting.** The user can explore the RDR and BAF data across samples in a local and global fashion.

*(P1) Plot the RDR and BAF values of bins for each sample sorted by genomic coordinates.* To facilitate local segmentation, the user can inspect RDR and BAF values ordered by genomic coordinates. This will enable the user to identify breakpoints along the genome.

*(P2) Simultaneously plot RDR and BAF values of bins for each sample.* To facilitate global segmentation, the user can inspect RDR and BAF values in a two-dimensional plot for each sample. This will enable the user to identify groups of bins distributed across the genome with similar RDR and BAF values across all samples.

*(P3) Indicate clustering of bins with the same set of colors in all plots.* To enable the user to view the current clustering, the tool should indicate cluster assignments of bins with colors. Specifically, the same set of colors is used in both the local plots (P1) as well as the global plots (P2).

*(P4) Show input data and cluster assignment of an individual bin.* The user can inspect the input data of an individual bin as well as its assigned cluster (if any).

**Filtering.** As we envision a tool that enables both local and global segmentation, the user can set filters in both a localized manner as well as a global manner.

*(F1) Show only bins that occur in a localized genomic range.* The user can restrict the shown bins to only those that occur within a user-specified linear genomic range via the local plots introduced in (P1).

*(F2) Show only bins that occur within a range of BAF and RDR values.* The user can specify a range of RDR and BAF values for each sample, restricting the tool to only show those bins whose RDR and BAF values occur within the specified ranges.

*(F3) Zooming and panning can be reset to a default state where all bins are shown.* This can be done on both the local and global plots.

*(F4) Show only bins that occur on an individual chromosome.* The user can specify an individual chromosome, restricting the tool to only show those bins that occur on the specified chromosome.

*(F5) Show only bins that are assigned to a specified set of clusters.* The user can specify one or more clusters, restricting the tool to only show those bins that are assigned to any of the specified clusters.

*(F6) The same set of bins should be shown in both the local and global plots for each sample.* To maintain visual consistency, it is important to ensure that the same set of bins is shown for each sample at all times. This is particularly important when altering filtering criteria via any of the aforementioned filtering tasks.

**Selecting.** Selection is an important step to enable the user to identify a group of bins and subsequently update their cluster assignments.

*(S1) A range of localized bins can be selected and deselected.* The user can add a range of bins to the current selection via the local plots introduced in (P1). Conversely, the user can remove bins from the current selection in a localized fashion.

*(S2) A set of bins with similar RDR and BAF values in one sample can be selected and deselected.* The user can add a set of bins with similar RDR and BAF values to the current selection via the global plots introduced in (P2). Moreover, the user can remove bins from the current selection using the same global plots.

*(S3) The current selection can be cleared.* The user can quickly clear the current selection (i.e. deselect all bins).

*(S4) The current selection of bins must be shown in all local and global plots.* The user can view the current selection in all plots across all samples.

**Clustering.** Next, the user can cluster the selected bins using the following functionality.

*(C1) The current selection of bins can be assigned to a new cluster.* Specifically, the tool should identify a new cluster index that has not been previously used and assign the bins to this cluster. If a subset of the selected bins were previously assigned to another cluster, they should be re-assigned to the new cluster.

*(C2) The current selection of bins can be merged into an existing cluster.* The user can select a previous cluster and assign the selected bins to this cluster. If a subset of selected bins were previously assigned to another cluster, they should be re-assigned to the cluster selected by the user.

*(C3) The cluster assignments of the selected bins can be cleared.* This functionality should reset the cluster assignments of the selection without assigning the selected bins to an existing cluster.

*(C4) All cluster assignments can be cleared.* The user can quickly clear cluster assignments of all bins.

*(C5) The current selection of bins can be erased.* To enable the user to remove outlier bins from consideration by downstream copy-number callers, the tool should provide functionality to erase the current selection.

*(C6) Any of the aforementioned clustering tasks can be undone.* To enable the user to recover from any mistakes during clustering (without starting over), the tool should maintain an undo stack.

*(C7) A log of all clustering operations is maintained.* To facilitate reproducibility, the tool should maintain an exportable log of all operations.

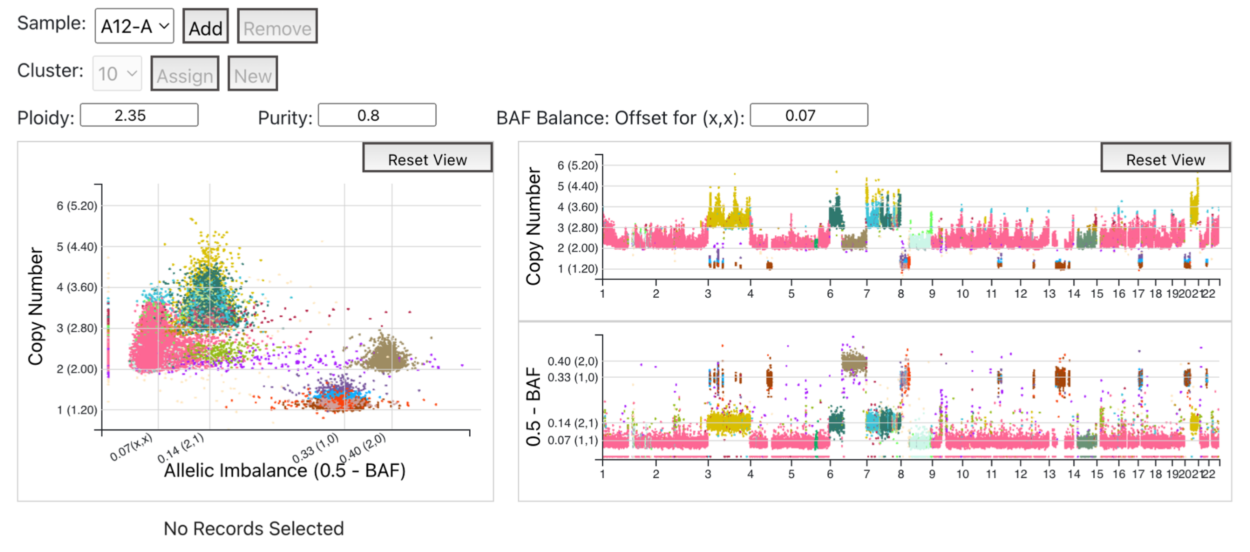

Fig A: **CNAViz allows the user to explore purity and ploidy values.** Tumor purity is the proportion of tumor cells in a sample, and the ploidy is the average number of copies. The user can vary the tumor purity and ploidy for each sample, and subsequently pick better parameter values for the downstream copy-number estimation process.

**Analytics.** The tool provides the user with feedback reflecting the current clustering.

*(A1) Show metrics assessing homogeneity and separation for each cluster.* A good clustering satisfies the following two criteria. First, bins within each cluster have similar RDR and BAF values per sample (i.e. homogeneity or cohesion). Second, distinct clusters are comprised of bins with distinct RDR and BAF values per sample (i.e.ṡeparation). Our tool should provide the user feedback of the current clustering regarding these two criteria. In particular, the tool should identify clusters that would benefit from further refinement.

*(A2) Show centroids of currently selected clusters.* To enable the user to visually assess the clustering, the tool can show cluster centroids in the global plots (P2). Only centroids for the currently selected clusters (F5) should be shown.

*(A3) Show driver gene locations.* To facilitate interpretation of CNAs, the tool should show driver gene locations in the linear plots (P1). The set of shown driver genes can be customized by the user (D6).

*(A4) Show gridlines illustrating purity and ploidy.* As picking the correct purity and ploidy values is a significant challenge in the downstream step of copy number calling, our tool should provide a grid outline for users to explore how different values of purity and ploidy broadly translate to copy number calls. Figure A illustrates the user interface.

**Automated Analytics.** Finally, CNAVIZ provides the user with a few automated functions.

*(AA1) Merge clusters.* Automates the ability to merge clusters with a button. The user provides RDR and BAF thresholds per sample; if the distance between two centroids is smaller than the thresholds in all samples, then the pair of clusters is merged. When a pair of clusters is merged, the union of the bins in both clusters is assigned to the largest starting cluster.

*(AA2) Split clusters.* Automates the ability to split clusters with a button. The user provides RDR and BAF thresholds per sample, and identifies a set of “From” and a set of “To” clusters. For a given bin in a “From” cluster, the distance from that bin to each “To” cluster centroid is calculated, and the minimum centroid is identified. If the RDR and BAF distance between this bin and the identified minimum centroid

is below the user-specified thresholds in each sample, then the bin in question is reassigned to the minimum distance centroid.

### B CNAVIZ Usage

We provide instruction on how to obtain CNAVIZ's input from BAM files, i.e. alignments of sequencing reads (Section B.1). Next, we detail how to convert a local segmentation obtained from ASCAT [1, 2] for further processing with CNAVIZ (Section B.2). We then describe how to obtain a global segmentation using HATCHet [3] (Section B.3). We also describe how an off-the-shelf GMMHMM approach can be used to obtain an initial clustering for CNAVIZ (Section B.4). We conclude by discussing how to perform CNA calling using CNAVIZ's output with HATCHet [3] (Section B.5). For further details we refer the reader to the tutorial available on `https://github.com/elkebir-group/cnaviz`.

#### B.1 Extracting RDR and BAF values from a BAM File

We provide the user with a script to use HATCHet to obtain RDR and BAF values per genomic bin when given a reference panel of germline SNPs as well as BAM files of one or more tumor samples and a matched normal sample. This script is published on the github repository (`https://github.com/elkebir-group/cnaviz/blob/master/docs/hatchet_rdrbaf.ini`).

#### B.2 Generating an Initial Clustering using ASCAT

The user can run ASCAT to generate an initial clustering. We provide an example script on how to perform the ASCAT clustering (`https://github.com/elkebir-group/cnaviz/blob/master/data/ascat/ASCAT_casasent.R`). After running ASCAT, to save files relevant to CNAVIZ we use the following R commands on the ASCAT output:

```
write.csv(ascat.output$segments)
write.csv(ascat.bc$SNPpos)
```

To reformat the ASCAT clustering into CNAVIZ input format, we provide the user with a published script as well (`https://github.com/elkebir-group/cnaviz/blob/master/data/ascat/ascat_outputs/ascat2cnaviz_input.py`).

#### B.3 Generating an Initial Clustering using HATCHet

The user can also run HATCHet to generate an initial clustering. We provide the user with scripts to obtain a HATCHet clustering on the github repository (`https://github.com/elkebir-group/cnaviz/blob/master/docs/hatchet_pre.ini`). These scripts require the user to input a BAM file containing tumor and normal samples, as well as a SNP reference panel. The HATCHet output can directly be imported by CNAVIZ.

#### B.4 Generating an Initial Clustering using GMMHMM

We explored default clustering solutions that framed the trade-offs between local and global segmentation solutions as an optimization problem. Local algorithms have the advantage of using the genomic locations of read bins. On the other hand, global algorithms use RDR and BAF information aggregated across the genome. Given both of these approaches, the problem is to find a function that solves the standard clustering problem while minimizing the number of segments created.

##### B.4.1 Gaussian Mixture Model-Hidden Markov Model

One possible solution to solving the copy number calling problem posed is to use a Gaussian Mixture Model. As shown in HATCHet, fitting a distribution to a set of RDR/BAF values has been shown to be an effective approach to obtain clusters which correspond to CNAs. We draw on this work to fit a Gaussian mixture model to the RDR and BAF values of each bin.

However, relying on RDR and BAF values alone is not as effective at capturing CNAs that are smaller in scale. To resolve this issue, we can use the position of a read bin in the genome to inform a hidden Markov model. Therefore, we describe a two-step process, where the first step leverages the Guassian mixture model (GMM), and the second step leverages the GMM emission and transition probabilities to inform the hidden Markov Model (HMM).

The means and covariance fit in the GMM are used to calculate the emission probability for each bit via the multivariate Gaussian probability density function. Using this emission matrix, and a uniform probability transition matrix, we construct an HMM and use the Viterbi algorithm to find the best path (in this case this translates to the best clustering assignment for each bin). We refer to this as the GMM-HMM approach. We use the GMM-HMM class in the widely used `hmmlearn` Python package.

##### B.4.2 Model Selection

Since there is no model selection built into the GMMHMM model, we perform model selection by using two metrics: the maximum likelihood score of a clustering assignment, and the silhouette score.

Figure B illustrates the model selection run on the A12 patient. On the $x$-axis we have a varying number of clusters. On the left $y$-axis we have the likelihood of each clustering solution, and on the right $y$-axis we have the silhouette score of each clustering solution. This was run with the scripts published on the github repository (`https://github.com/elkebir-group/cnaviz/tree/master/initial_clustering`).

We used the following command to run the GMM+HMM on A12:

```
python model.py --input_file a12.tsv
   --num_restarts 2 --num_clusters_min 2
   --num_clusters_max 22 --num_clusters_step 1
   --num_processes 6 --output_folder a12_gc
```

The script will run GMMHMM $\lceil \frac{\texttt{num\_clusters\_max} - \texttt{num\_clusters\_min} + 1}{\texttt{num\_clusters\_step}} \rceil \times \texttt{num\_restarts}$ times, find the likelihood of the model given the data and the silhouette score, store the results in a format that allows the user to import the clustering into CNAVIZ, and outputs a plot that shows the user the scores to determine which number of clusters is the best in terms of these two scores. Figures B and C illustrate the 3-dimensional model which includes RDR, BAF and genomic positions of each bin.

By evaluating Figure B, we select the solution containing 7 clusters, which has the best tradeoff with a high likelihood and relatively high silhouette score. For comparison, Figure C shows two solutions: the first solution contains 7 clusters, as this has the best tradeoff between likelihood and silhouette score. The second solution contains 22 clusters and has a high likelihood but lower silhouette score and more clusters.

In Figure C we can immediately observe that there are some clear mistakes in the left hand column (clustering solution with 7 clusters). In particular, the dark blue and cyan blue share the same RDR and BAF values across all samples, and should be one cluster. Additionally, the cyan cluster is split in the BAF values in chromosome 6 across both samples, and so should be split into two clusters. On the other hand, in the right hand column (clustering solution with 22 clusters) of Figure C, we can observe that several spurious clusters should instead be one cluster as they share an RDR and BAF value across both samples. Additionally, the green cluster should clearly have bene split in the 2D scatterplot in panel (d).

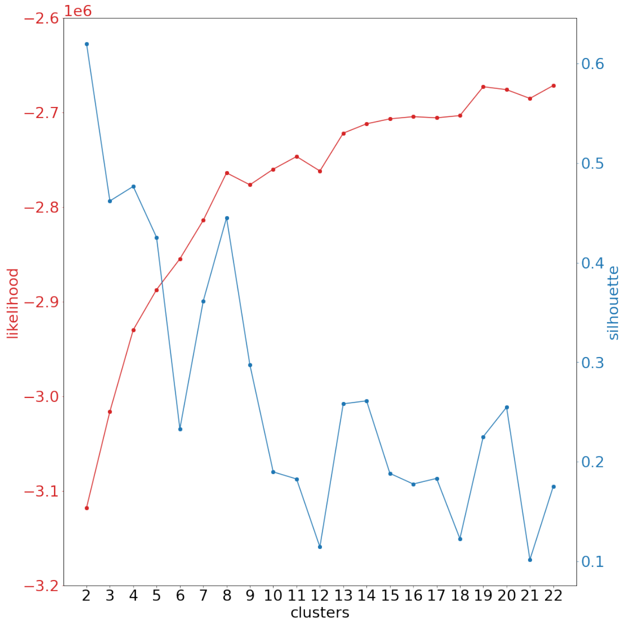

Fig B: **Likelihood versus Silhouette score over cluster numbers plot to compare results over clusterings for patient A12.** Each red point represents a clustering's likelihood score, and the blue line indicates the silhouette score for the best likelihood score from each clustering number.

Subsequently, we can observe that the ability of the initial clustering to produce a segmentation is quite limited. Solving this optimization problem where we balance the genomic coordinates and the RDR and BAF values, as well as perform a parameter sweep for purity and ploidy and perform model selection to identify the best number of clusters, is extremely difficult, and almost always makes mistakes. CNAVIZ was designed specifically for this use case where heuristics may fail to identify mistakes, but where they are clear by visual inspection.

##### B.5 Downstream CNA Calling after CNAVIZ

After manual segmentation with CNAVIZ, the user is free to leverage any existing copy number calling methods they wish to perform this downstream process. We provide the user with a script to perform this copy number calling process with HATCHet on the github repository (`https://github.com/elkebir-group/cnaviz/blob/master/docs/hatchet_post.ini`).

#### C Results

##### C.1 Simulated Data

We used CNAVIZ to perform manual clustering on the published dataset `n2_s4669` (no whole genome duplications) simulated with MASCoTE [3]. We use the simulated version of `n2_s4669` which contains 4 bulk DNA sequencing samples with 2 tumor clones. This dataset was published with the RDR and BAF values already calculated.

Since `n2_s4669` was published with a HATCHet clustering performed by the authors, we did not re-run this step. To run ASCAT, we first converted the simulated data to the required input format (script published here: `https://github.com/elkebir-group/cnaviz/blob/master/data/ascat/`

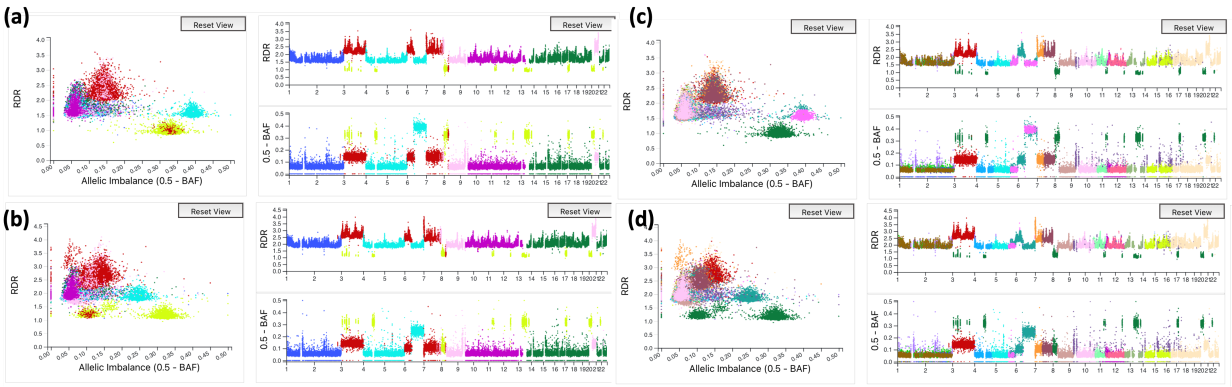

Fig C: **Shows two clustering solutions on patient A12 in the CNAViz user interface.** The left column shows a clustering solution containing 7 clusters, with a silhouette score of approximately 0.37. Sample A is shown in panel (a), and Sample B is shown in panel (b). A second clustering solution containing 22 clusters with a silhouette score of approximately 0.185. Sample A is shown in panel (c) and Sample B is shown in panel (d).

`ascat_inputs/ascat_input.py`). Then, we ran ASCAT in the `aspcf` mode. We also ran the `multipaspcf` mode, but as this version's segmentations produced even more clusters than the `aspcf` mode, we used the `aspcf` results for ease of manual editing. We ran ASCAT in the `aspcf` mode with manually calculated ASCAT input values of rho and psi for each sample in the simulated `n2_s4669` dataset: `rho_manual`=c(0.949758, 0.949751, 0.799842), `psi_manual`=c(1.971723, 1.951668, 0.923111)), obtained from ground truth. The script used to calculate these values of rho and psi (purity and ploidy) is also published on GitHub (`https://github.com/elkebir-group/cnaviz/blob/master/data/ascat/ascat_inputs/calc_rho_psi.py`). The script used to run ASCAT is published on the GitHub repository `https://github.com/elkebir-group/cnaviz/blob/master/data/ascat/ASCAT_casasent.R`. To reformat the ASCAT clustering into CNAViz input format, we provide the user with a published script as well (`https://github.com/elkebir-group/cnaviz/blob/master/data/ascat/ascat_outputs/ascat2cnaviz_input.py`).

In order to generate copy number calls after manual editing with CNAViz, we used the copy number call solver built into HATCHet, but the user can use any copy number caller of their choice as CNAViz outputs standard information about each bin. The scripts to generate copy number calls from HATCHet are provided on the github repository as well (`https://github.com/elkebir-group/cnaviz/blob/master/docs/hatchet_post.ini`), with a more detailed data preparation tutorial available as well (`https://github.com/elkebir-group/cnaviz`).

#### C.2 Real Data

We used patients P5 P6 and P10 from the Casasent et al. 2018 [4] dataset, which were also used in the HATCHet paper, and for which results are published.

We reran HATCHet v0.4.7 on this `n2_s4660` dataset with default settings. The scripts used to obtain a HATCHet clustering are published on the github repository (`https://github.com/elkebir-group/cnaviz/blob/master/docs/hatchet_pre.ini`).

We ran ASCAT in the `aspcf` mode, which we describe above, with all the same input scripts. The calculated rho and psi values (which can also be found in the script used to run ASCAT) for each sample are as follows:

- P5: rho (0.680271, 0.431884), psi (4.178528, 3.346920)
- P6: rho (0.768595, 0.820807), psi (1.599508, 1.445370)
- P10: rho (0.961155, 0.844639), psi (1.567808, 1.644032)

In order to generate copy number calls after manual editing with CNAVIZ, we used the same scripts as mentioned in the simulated data section above.
